# Nanopore-only assembly and methylation profiling of *Arabidopsis thaliana* across chemistries, basecalling models and assembly workflows

**DOI:** 10.1101/2024.11.13.623402

**Authors:** Samuel Martin, Ned Peel, Darren Heavens, Yuxuan Lan, Naomi Irish, Samuel T. Horsfield, Michael Giolai, Matthew D. Clark, Richard M. Leggett

## Abstract

Over the past decade, nanopore sequencing has progressed rapidly through advances in flow cell chemistry, library preparation, basecalling algorithms and base-modification detection. These developments have increased both throughput and read accuracy, making nanopore-only sequencing increasingly practical for eukaryotic genome assembly and direct detection of DNA methylation. Here, we present nanopore sequencing data, genome assemblies, and methylation analysis for the Columbia (Col-0) accession of the model plant *Arabidopsis thaliana* and assess how successive improvements in the nanopore sequencing platform have affected read accuracy and assembly quality. Data from earlier R9.4.1 and R10.4 flow cells were sufficient to generate nanopore-only multi-megabase and chromosome-arm scale assemblies, with higher reference-alignment identity achieved with the improved R10.4 chemistry. Super-accuracy rebasecalling increased read quality in both datasets and markedly improved assembly identity and gene completeness. Methylation calls obtained directly from nanopore signal were strongly concordant with a published bisulphite sequencing dataset at shared covered sites, with Pearson’s r values of 0.93 and 0.96 for the R9.4.1 and R10.4 datasets, respectively. To assess current nanopore assembly performance, we additionally analysed 14.3 Gb R10.4.1 simplex reads, basecalled with a super-accuracy model and assembled using hifiasm, a contemporary long-read de novo assembler. This produced a five-contig nuclear assembly of 134.93 Mb, with each contig corresponding to one of the five *Arabidopsis* nuclear chromosomes. The assembly recovered 99.9% complete BUSCO genes and showed 99.98% one-to-one alignment identity to TAIR12, with three chromosomes assembled telomere-to-telomere. These results provide an empirical example of how nanopore-only plant genome assembly has improved across successive workflows and demonstrate the value of nanopore data for simultaneous genome assembly and methylation analysis.

## Introduction

Nanopore sequencing has transformed genome analysis by enabling direct, single-molecule sequencing of native DNA in real time. In contrast to short-read platforms, Oxford Nanopore Technologies (ONT) platforms can generate reads ranging from several kilobases to hundreds of kilobases, with ultra-high molecular weight DNA preparation extending read lengths into the megabase range (Cahyani et al., 2025). Long reads more generally are useful for resolving repetitive regions, structural variants, haplotypes, and other genomic features that are difficult to reconstruct from short-read data alone. Since the release of the portable MinION platform through the MinION Access Programme (MAP) in 2014, successive improvements in pore design, sequencing chemistry, library preparation, and basecalling algorithms have substantially increased both yield and accuracy of sequenced molecules (Leggett and Clark, 2017; Sereika et al., 2022). These advances have expanded ONT sequencing from early applications in microbial genomics to increasingly complex eukaryotic genomes.

*Arabidopsis thaliana* is a well-suited model system for assessing plant genome assembly methods. It is diploid and has a compact nuclear genome comprising five chromosomes, approximately 135 Mbp in size, with repetitive regions, albeit relatively low total repeat content compared with many other plant genomes (Quesneville, 2020; Lee and Kim, 2014). *A. thaliana* is a heavily used genetic model system, and was the first plant genome to be sequenced and remains one of the best-supported plant reference systems, with extensive genomic, transcriptomic, proteomic, and epigenomic resources (The Arabidopsis Genome Initiative, 2000). Its Columbia accession, Col-0, is the basis of widely used reference assemblies, including TAIR10 and the recently released TAIR12 reference genome (Lamesch et al., 2012; Reiser et al., 2026). Beyond Col-0, the 1001 Genomes Project has characterised genetic variation across 1,135 *A. thaliana* accessions, primarily using short-read sequencing (1001 Genomes Consortium, 2016). Complementary long-read assemblies could extend this resource by improving the resolution of structural variation and repetitive regions. Together, these extensive genomic resources make *A. thaliana* a useful benchmark for evaluating how sequencing and analysis workflows affect assembly contiguity, consensus accuracy, and gene completeness, with potential applications to other plant species that have less well-developed genomic resources.

Previous plant genome assembly studies have demonstrated the value of long nanopore reads, although many high-quality assemblies have relied on short-read polishing, very deep sequencing, ultra-long reads, or additional scaffolding technologies. A high-contiguity assembly of *A. thaliana* KBS-Mac-74 was previously generated from nanopore data and polished with Illumina reads (Michael et al., 2018), while hybrid approaches using long reads to scaffold short-read assemblies have been applied to other plant genomes (Perumal et al., 2020). Subsequent studies have used deep ultra-long nanopore sequencing to resolve gaps in the TAIR10 reference, and combined nanopore, PacBio HiFi, and chromosome-conformation data has been used to generate near-complete *A. thaliana* assemblies (Naish et al., 2021; Wang et al., 2022). These advances have contributed to the improved *Arabidopsis* reference landscape, culminating in community resources such as TAIR12, which incorporates long-read-enabled genome improvements and updated annotation (Reiser et al., 2026). Continued and frequent improvements in nanopore sequencing accuracy and long-read assembly methods make it useful to reassess nanopore-only assembly performance with current sequencing and analysis workflows.

Nanopore sequencing can also provide information about modified bases because DNA modifications alter the raw nanopore signal. This allows DNA methylation to be detected from the same sequencing data used for assembly, without bisulphite conversion or a separate methylation-specific library preparation (Simpson et al., 2017). Where raw signal data have been retained, they can also be reprocessed using updated modification-detection models, enabling methylation analysis without resequencing. Furthermore, long nanopore reads can link methylation calls to individual haplotypes, enabling allele-specific methylation analysis (Akbari et al., 2021). Bisulphite sequencing remains a widely used approach for measuring cytosine methylation, but it requires chemical conversion of DNA and usually short-read library preparation; bisulphite treatment and subsequent amplification can introduce DNA degradation and library-preparation biases (Olova et al., 2018). Nanopore-based methylation detection is therefore attractive for studies where long-range sequence context and methylation information are both useful, although methylation calls depend on the basecalling and modification-detection models used.

Here, we use *A. thaliana* Col-0 as a practical benchmark to assess progress in nanopore-only plant genome assembly and CpG methylation analysis across three successive nanopore datasets. We compare datasets generated using earlier R9.4.1 and R10.4 nanopore flow cells and assess how basecalling and chemistry improvements affected read accuracy, assembly contiguity, and consensus quality using several nanopore assembly and polishing workflows. We also compare CpG methylation calls obtained directly from nanopore data with a previously published bisulphite sequencing dataset. Finally, to assess current nanopore assembly performance, we analyse a dataset generated using the latest (at the time of writing) R10.4.1 chemistry and processed with recent basecalling and long-read assembly methods. Together, these analyses provide a practical benchmark for nanopore-only genome assembly and methylation analysis in a small plant genome and illustrate how successive improvements in nanopore sequencing and analysis software have increased the quality of assemblies that can be generated from modest sequence yield.

## Materials and Methods

### Plant material and DNA extraction

*Arabidopsis thaliana* Col-0 plants were grown for 10 days after germination and then kept in darkness for 48 h before harvest to reduce starch accumulation, secondary metabolites, and the relative contribution of plastid DNA. Harvested material was stored at −80°C before extraction. Leaf material was ground to a fine powder in liquid nitrogen using a mortar and pestle. DNA was extracted using the Illustra Nucleon Phytopure Genomic DNA Extraction Kit (Cytiva, Sheffield, UK), with Reagent 1 and Reagent 2 prewarmed to 65°C before extraction to improve DNA molecular weight.

Ground tissue (0.1 g) was added to 600 µl of Reagent 1 in a 2 ml Eppendorf LoBind tube and mixed thoroughly before 200 µl of Reagent 2 was added. The tube was incubated in a preheated Eppendorf ThermoMixer C at 500 rpm for 10 min at 65°C, then placed on ice for 20 min. Ice-cold chloroform (500 µl) and Nucleon Phytopure DNA extraction resin (100 µl) were added, and the tube was mixed by inversion on a lab rotator at 30 rpm for 10 min at room temperature. The tube was centrifuged for 10 min at 1,300 × g at 4°C. The upper aqueous phase was transferred to a fresh tube, mixed with an equal volume of ice-cold isopropanol, inverted five times and centrifuged for 5 min at 4,000 × g at 4°C. The supernatant was discarded, the pellet was washed twice with 70% ethanol, and DNA was dissolved in 50 µl DNase-free water overnight at 4°C.

DNA concentration was measured using the Qubit BR assay (Life Technologies, Loughborough, UK). DNA molecule length was assessed using an Agilent Femto Pulse system (Agilent, Cheadle, UK).

### Nanopore library preparation and sequencing

Nanopore libraries were constructed using ONT sequencing kits according to the manufacturer’s instructions, except that Kapa Pure Beads (Roche, UK) were used for clean-up steps instead of AMPure XP beads and no fragmentation step was applied. For each sequencing run, 1 µg of DNA was used as input for library construction.

The three sequencing datasets are referred to as ATRun1, ATRun2, and ATRun3. ATRun1 was prepared using the SQK-LSK109 kit, ATRun2 using the SQK-Q20EA kit, and ATRun3 using the SQK-LSK114 kit with the supplied Long Fragment Buffer to help remove library molecules <3kbp after adapter ligation. Each library was sequenced for 72 h. ATRun1 and ATRun2 were sequenced on individual R9.4.1 and R10.4 flow cells, respectively, using a GridION Mk1. ATRun3 was sequenced on a single FLO-PRO114M R10.4.1 flow cell using a PromethION P2 Solo. Run statistics and basecalling models are summarised in Table 1.

**Table 1:** Read statistics for nanopore datasets and basecalling models.

| **Run** | **Flow cell** | **Kit** | **Basecalling model** | **Total reads** | **Total bases (Gb)** | **Read N50 (kb)** | **Mean read length (kb)** | **Mean read Q score** | **Approx. genome coverage** |
|---|---|---|---|---|---|---|---|---|---|
| ATRun1 | R9.4.1 | SQK-LSK109 | HAC | 258,885 | 2.86 | 35.00 | 11.04 | 9.92 | 21× |
| ATRun1 | R9.4.1 | SQK-LSK109 | SUP | 258,885 | 2.91 | 35.22 | 11.22 | 13.65 | 22× |
| ATRun2 | R10.4 | SQK-Q20EA | HAC | 645,585 | 12.65 | 33.24 | 19.60 | 12.37 | 94× |
| ATRun2 | R10.4 | SQK-Q20EA | SUP | 645,585 | 12.78 | 33.56 | 19.79 | 14.36 | 95× |
| ATRun3 | R10.4.1 | SQK-LSK114 | HAC | 705,601 | 14.11 | 37.34 | 20.00 | 14.72 | 105× |
| ATRun3 | R10.4.1 | SQK-LSK114 | SUP | 706,072 | 14.30 | 37.54 | 20.25 | 19.64 | 106× |

Note: HAC, high-accuracy basecalling; SUP, super-accuracy basecalling. Approximate genome coverage was calculated using an assumed nuclear genome size of 135 Mb and includes any organellar reads present.

### Basecalling, read filtering, and read-level Q score calculation

PycoQC (Leger and Leonardi, 2019) was used to inspect read-level statistics for all sequencing runs. For each read, a read-level Q score summarising the per-base quality scores was calculated by converting each base Q score to an error probability, averaging error probabilities across the read, and converting the mean error probability back to a Phred-scaled Q score. Mean read Q scores reported in Table 1 are the mean of these read-level Q scores.

For ATRun1, reads with a mean Q score below 7 (corresponding to approximately 80% predicted basecalling accuracy) were removed using `QScore_filter.py`, and remaining reads were trimmed for adapter sequence using Porechop (Wick et al., 2017) before assembly. ATRun2 analyses used reads that passed MinKNOW quality control, followed by adapter trimming with Porechop.

FAST5 files from ATRun1 and ATRun2 were rebasecalled with Guppy using SUP models: Guppy v5.0.7 for ATRun1 and Guppy v5.0.16 for ATRun2. Reads passing Guppy quality control were trimmed for adapter sequence using Porechop. ATRun3 generated POD5 files rather than FAST5 files. ATRun3 raw signal data were basecalled with a HAC model (dna_r10.4.1_e8.2_400bps_hac@v4.3.0), and then rebasecalled using Dorado v0.8.3 with a SUP model (dna_r10.4.1_e8.2_400bps_sup@v5.0.0) for assembly. Dorado performed adapter trimming during basecalling. No additional read-quality or read-length filtering was applied to ATRun3, and all reads output by SUP basecalling were used for assembly. All analyses used simplex reads, with individual DNA strands basecalled independently. No duplex consensus reads were used.

### Alignment-based read accuracy assessment

Reads from each sequencing run were aligned to the TAIR10 *A. thaliana* reference genome using Minimap2 (Li, 2018). As the sequenced material was also Col-0, alignment mismatches were expected to predominantly reflect sequencing errors rather than genetic differences, allowing reference alignments to be used to estimate read accuracy. The `-c` option was used to request base-level alignment output and `--cs` was used to generate the difference string used for parsing alignments. Only reads with alignments covering at least 90% of the read length were retained for accuracy assessment. Where a read had multiple alignments meeting this criterion, the alignment with the greatest alignment length was used. Alignment-based read accuracy was calculated using the BLAST identity convention, defined as the number of exact matches divided by the alignment length. The scripts used to perform alignments, parse identity values, and plot read accuracy distributions are available at https://github.com/SR-Martin/RQbRA.

### Assembly and polishing of ATRun1 and ATRun2

Following read quality filtering and adapter trimming, ATRun1 and ATRun2 reads were assembled using Canu v1.8 (Koren et al., 2017), Flye v2.8 (Kolmogorov et al., 2019), Miniasm v1.0 (Li, 2016), and wtdbg2 v2.5 (Ruan and Li, 2020). Assemblies produced by Canu, Flye, Miniasm, and wtdbg2 were polished with Racon v1.3.1 (Vaser et al., 2017) and then polished using Nanopolish v0.13.2 (Loman et al., 2015) with the corresponding FAST5 signal data. ATRun1 and ATRun2 read sets were also assembled using the ONT Pomoxis pipeline v0.2.2 and polished using Medaka v0.6.2, as recommended for that workflow.

Separate polishing experiments were performed using ATRun1 assemblies distinct from those reported in Table 2. Table S1 compares assemblies before and after polishing. Table S2 compares one versus four rounds of Racon polishing of a Flye assembly; both treatments were followed by a final Nanopolish step.

### ATRun3 assembly and curation

The ATRun3 SUP reads were assembled using hifiasm v0.25.0-r726 (Cheng et al., 2026) in ONT mode with the following command:

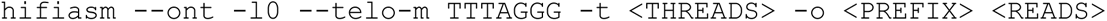

Coverage information from hifiasm GFA rd:i tags was used to identify high-coverage assembly contigs. A threshold of rd:i ≥ 60 was applied to generate a coverage-filtered assembly. This threshold was chosen after inspection of the rd:i distribution because it separated the main high-coverage contigs from lower-coverage contigs. High-coverage contigs were classified using BLAST+ v2.10.0 (Camacho et al., 2009) against NCBI core nt database version 2026-05-04. Contigs classified as organellar were excluded from the final nuclear assembly. The remaining five high-coverage contigs were retained as the curated nuclear assembly.

### Assembly evaluation

Assembly contiguity was summarised using a custom script (`get_stats.py`) to calculate the number of contigs, total assembly length, and N50. ATRun1 and ATRun2 assemblies were compared with the TAIR10 reference genome using dnadiff from MUMmer v3.23 (Kurtz et al., 2004). Gene completeness was assessed using BUSCO v4.0.2 (Seppey et al., 2019) with the brassicales_odb10 dataset. Structural correspondence between assemblies and TAIR10 was assessed by aligning assemblies to the reference with Minimap2 and visualising alignments with Alvis (Martin and Leggett, 2021).

ATRun3 assemblies were evaluated separately against the TAIR12 reference genome. BUSCO completeness was assessed using BUSCO v5.8.3 with the brassicales_odb10 dataset. Consensus quality value (QV), a Phred-scaled estimate of consensus accuracy, was estimated using yak v0.1-r69 (https://github.com/lh3/yak). A k-mer database was constructed from the same Dorado SUP reads used for assembly using `yak count -b37 -t 4`, and assemblies were evaluated against this database using `yak qv -t 4`. Adjusted QV estimates were reported. These estimates quantify consensus accuracy inferred from read k-mer support. Because the same reads were used for assembly and evaluation, they may be affected by shared systematic errors. Reference-based comparison was performed with MUMmer v4.0.1 using nucmer and dnadiff against TAIR12. Whole-contig alignment visualisations were generated by aligning the curated five-contig nuclear assembly to TAIR12 using Minimap2 and visualising the resulting alignment file with Alvis. These alignments were used to assess the correspondence between individual assembly contigs and the five *A. thaliana* nuclear chromosomes. Alignments shorter than 1% of the corresponding reference sequence were excluded from visualisations.

### Nanopore methylation calling and comparison with bisulphite sequencing

Public Illumina bisulphite sequencing data for *A. thaliana* Col-0 leaf tissue were obtained from NCBI GEO accession GSE43857, sample GSM1085222. Only cytosine methylation calls at CpG sites were used for comparison with nanopore methylation calls.

Nanopore methylation detection was performed by calling modified bases from FAST5 data using Guppy with alignments to the TAIR10 reference genome. For ATRun1, Guppy v5.0.16 was used with the model dna_r9.4.1_450bps_modbases_5mc_hac. For ATRun2, Guppy v6.5.7 was used with the model dna_r10.4_e8.1_modbases_5mc_cg_sup. Modified-base BAM files were converted to BED format using modbam2bed v0.10:

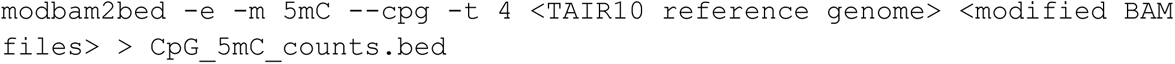

BED files were converted using a custom script (`bed_to_methylcall.py`) to match the coordinate and methylation-call format of the bisulphite sequencing dataset. Only chromosomal sites were retained. Using methylKit (Akalin et al., 2012), sites with coverage below 10 reads for nanopore data or 8 reads for bisulphite data were removed before strand merging. For each dataset, counts from complementary cytosines within a CpG were combined using `methylKit::unite()`. Each nanopore dataset was matched separately to the bisulphite dataset, retaining CpG sites represented in both datasets. Pearson’s correlation coefficient (r) between per-site methylation proportions was calculated using `methylKit::getCorrelation()` with default parameters.

### Chromosome-scale methylation profile

Chromosome 1 methylation profiles were generated using a JupyterLab notebook (`plots.ipynb`). Chromosome 1 was divided into non-overlapping 2.5 kbp bins. For each bin, the proportion of methylated CpG sites was calculated. A CpG site was classified as methylated if more than 50% of reads covering that site contained a 5mC call. Only sites passing the coverage thresholds described above were included.

### Gene-region methylation profile

Gene-region methylation profiles were generated using a custom script (`write_gene_meth_hists.py`). Two input gene sets were defined using leaf expression data from the Arabidopsis tissue atlas RNA-seq baseline dataset of Mergner et al. (2020), accessed through EMBL-EBI Expression Atlas (Moreno et al., 2022): 1,001 highly expressed genes and 5,552 genes with no detectable expression in leaf tissue. The number of genes represented in each region depended on the availability of the corresponding annotation and methylation calls. Gene coordinates were obtained from the TAIR10 annotation. Gene-region methylation profiles were calculated separately for 1-kb upstream regions, 5′ UTRs, exons, introns, 3′ UTRs, and 1-kb downstream regions. Each region type was represented by 100 normalised bins. Within each expression group, bin values were calculated by averaging the methylation proportions of contributing CpG sites pooled across genes.

### Use of artificial intelligence tools

ChatGPT (OpenAI) was used to assist with manuscript editing, including improving clarity, structure, and consistency of wording. It was also used to assist with drafting and checking small pieces of analysis code. All AI-assisted text and code were reviewed, edited, and verified by the authors, and all analyses and interpretations reported in the manuscript are the authors’ own.

## Results

We analysed three nanopore sequencing datasets generated from *A. thaliana* Col-0: ATRun1, ATRun2, and ATRun3. These datasets represent successive nanopore chemistries and analysis workflows: ATRun1 used R9.4.1 chemistry, ATRun2 used Q20+/R10.4 chemistry, and ATRun3 used R10.4.1 chemistry. ATRun1 and ATRun2 were used to compare earlier nanopore chemistries, basecalling models, assembly workflows, and CpG methylation detection. ATRun3 was used to assess current nanopore-only assembly performance using recent basecalling and long-read assembly methods. Sequencing and read statistics are summarised in Table 1.

### Sequencing yield and quality increased across nanopore datasets

Sequencing yield increased across the three successive datasets. ATRun1 produced 2.86–2.91 Gbp of data depending on the basecalling model, corresponding to approximately 21–22× genome coverage (Table 1). ATRun2 produced 12.65–12.78 Gbp, corresponding to approximately 94–95× coverage, while ATRun3 produced 14.11–14.30 Gbp, corresponding to approximately 105–106× coverage. Mean read length increased from approximately 11 kbp for ATRun1 to approximately 20 kbp for ATRun2 and ATRun3. Mean read Q scores were higher after SUP basecalling than HAC basecalling for all three datasets. The longest reads in ATRun1, ATRun2, and ATRun3 were 173 kbp, 271 kbp, and 1.46 Mbp, respectively. Reads longer than 50 kbp contributed approximately 5.7×, 21.0×, and 30.4× genome-equivalent coverage, respectively (Table S3).

### R9.4.1 data and earlier methods generated multi-megabase and chromosome-arm-scale assemblies

Data from a single R9.4.1 flow cell (ATRun1) were sufficient to generate multi-megabase *A. thaliana* assemblies using several long-read assemblers. The HAC read set contained 258,885 reads with a total length of 2.86 Gbp, corresponding to approximately 21× genome coverage, and a read N50 of 35.00 kbp (Table 1). After removal of reads with mean Q score below 7 and adapter trimming, 243,084 reads remained, with a total length of 2.80 Gbp, mean read length of 11.5 kbp, and read N50 of 35.3 kbp.

Among HAC assemblies, Flye produced the most contiguous R9.4.1 assembly, with 53 contigs, an N50 of 14.85 Mbp and 99.83% of TAIR10 bases covered (Table 2). However, consensus accuracy and gene completeness were limited for HAC assemblies: average alignment identity ranged from 98.70% to 99.23%, and BUSCO completeness ranged from 51.0% to 72.4% complete BUSCOs depending on assembler. The Pomoxis/Medaka workflow recovered the highest BUSCO completeness among HAC assemblies.

**Table 2:** Effect of rebasecalling on ATRun1 assembly quality. BUSCO values show complete (C), fragmented (F), and missing (M) BUSCOs using the brassicales_odb10 dataset.

| **Basecalling model** | **Assembler** | **Contigs** | **Total length (Mb)** | **N50 (Mb)** | **TAIR10 bases covered (%)** | **Average alignment identity (%)** | **BUSCO** |
|---|---|---|---|---|---|---|---|
| HAC | Canu | 124 | 121.17 | 10.76 | 99.74 | 98.88 | C:56.4%, F:9.4%, M:34.2% |
| HAC | Flye | 53 | 120.74 | 14.85 | 99.83 | 98.92 | C:57.5%, F:9.8%, M:32.7% |
| HAC | Miniasm | 42 | 118.85 | 8.38 | 99.66 | 98.70 | C:51.0%, F:9.2%, M:39.8% |
| HAC | Pomoxis | 82 | 123.71 | 10.07 | 99.71 | 99.23 | C:72.4%, F:7.7%, M:19.9% |
| HAC | Wtdbg2 | 286 | 127.03 | 7.49 | 99.66 | 98.74 | C:53.2%, F:10.7%, M:36.1% |
| SUP | Canu | 92 | 123.96 | 9.49 | 99.80 | 99.86 | C:97.3%, F:0.6%, M:2.1% |
| SUP | Flye | 51 | 119.86 | 14.64 | 99.83 | 99.86 | C:97.4%, F:0.6%, M:2.0% |
| SUP | Miniasm | 62 | 121.23 | 5.52 | 99.55 | 99.62 | C:97.2%, F:0.6%, M:2.2% |
| SUP | Pomoxis | 153 | 130.74 | 6.52 | 99.79 | 99.80 | C:96.3%, F:0.9%, M:2.8% |
| SUP | Wtdbg2 | 192 | 130.04 | 12.34 | 97.90 | 99.86 | C:97.1%, F:0.5%, M:2.4% |

Additional polishing produced modest, assembler-dependent changes in R9.4.1 assembly metrics. Assemblies produced by Canu, Flye, Miniasm and wtdbg2 were polished with Racon and Nanopolish, while Pomoxis assemblies were polished with Medaka. Polishing generally improved either consensus identity or contiguity, although not every metric improved for every assembler (Table S1). We also tested additional polishing of the Flye assembly by applying multiple rounds of Racon, following Vaser et al. (2017), before a final Nanopolish step. We found that this resulted in a slightly smaller assembly, but also a small loss in accuracy (Table S2).

### The super accuracy basecaller improves read accuracy and R9.4.1 assembly accuracy

Rebasecalling ATRun1 raw signal data with the SUP model increased the mean read Q score from 9.92 to 13.65, while read length distributions remained broadly similar (Table 1; Fig 1A). This improvement in read-level quality translated into higher alignment identity and BUSCO completeness across all evaluated assemblers. Average alignment identity increased from 98.70– 99.23% for HAC assemblies to 99.62–99.86% for SUP assemblies (Table 2). BUSCO completeness also increased markedly, from 51.0–72.4% complete BUSCOs in HAC assemblies to 96.3–97.4% in SUP assemblies (Fig 1B). Improvements in contiguity were less consistent: contig counts and N50 values did not improve for all assemblers.

**Fig 1.**
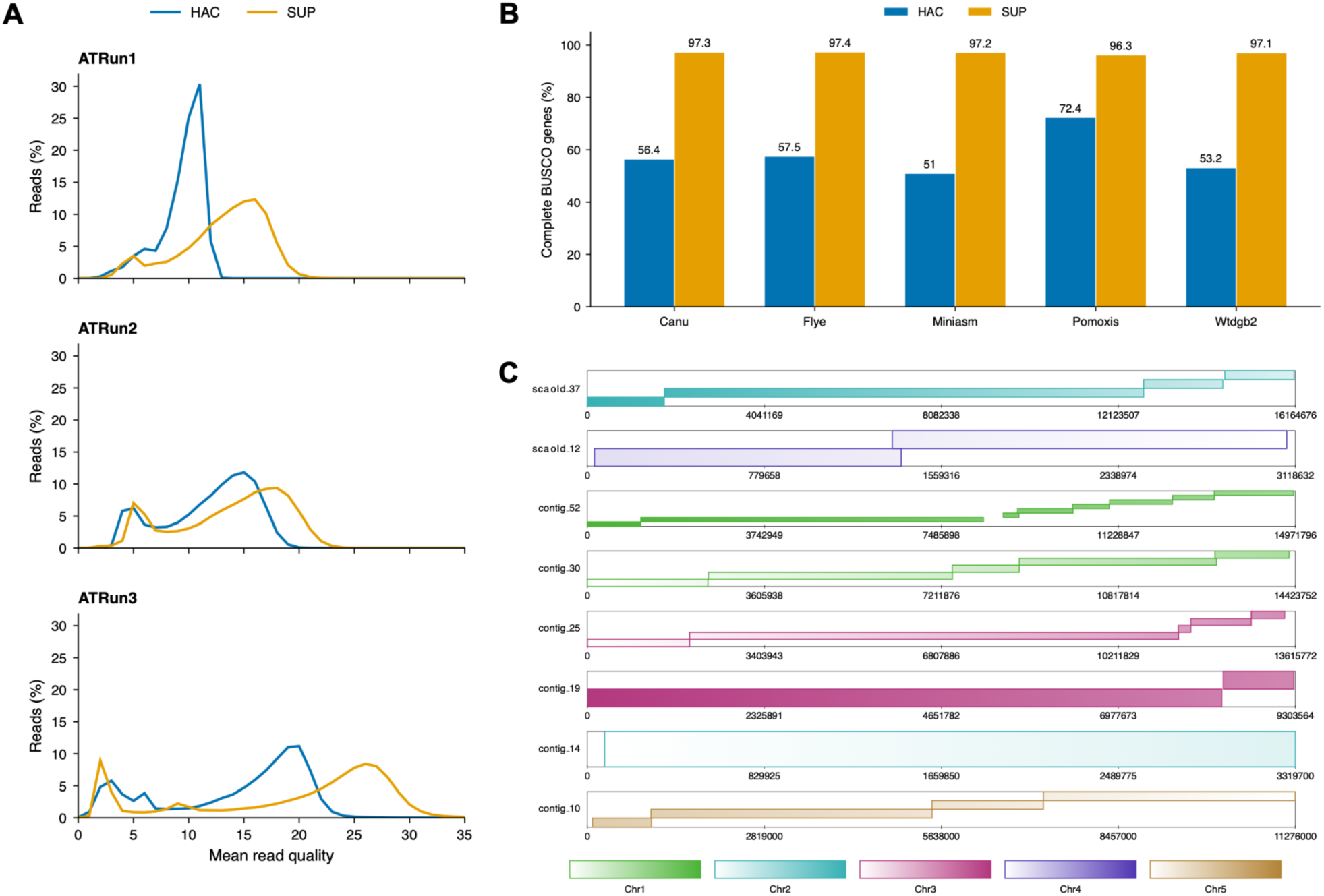
Read quality and assembly characteristics of nanopore datasets. **A.** Read-level Q score distributions for each run, shown for HAC and SUP basecalling models. **B.** Percentage of complete BUSCO genes found in assemblies of ATRun1, for both HAC and SUP basecalling. **C.** Visualisation of genome alignment of representative contigs from the assembly of ATRun1(SUP) by Flye against the TAIR10 reference genome. Each block represents a sequence in the assembly, the coloured blocks inside represent alignments to the reference sequences, shaded to indicate position and orientation. Small alignments (<1% of the reference sequence) have been filtered out.

Structural comparison of the ATRun1 SUP Flye assembly against TAIR10 showed that the longest assembly sequences corresponded largely to chromosome arms rather than complete chromosomes (Fig 1C). Visualisation of Minimap2 alignments for sequences longer than 1 Mbp showed, for example, that contig_30 and contig_52 aligned to opposite arms of chromosome 1, with contig_52 in the reverse orientation. Similarly, scaffold_37 aligned to the long arm of chromosome 2, while contig_14 aligned to the short arm. These alignments indicate that the R9.4.1 SUP Flye assembly reached chromosome-arm scale but did not resolve all chromosomes as single contigs.

### Q20+/R10.4 chemistry further improved read and assembly accuracy

ATRun2, generated using SQK-Q20EA chemistry and an R10.4 flow cell, produced substantially greater sequence yield than ATRun1 and higher alignment-based read accuracy (Table 3; Fig 2). Assembly of ATRun2 SUP reads using the same assembly and polishing workflows as ATRun1 produced average alignment identities of 99.84–99.96% across assemblers. Canu, Flye, and Pomoxis each covered more than 99.6% of TAIR10 bases, with N50 values of 12.62 Mbp, 13.59 Mbp, and 10.89 Mbp, respectively. However, assemblies remained fragmented relative to the five nuclear chromosomes, with 49–195 contigs for Canu, Flye, Miniasm, and Pomoxis. Polishing failed for the wtdbg2 assembly, which had lower reference coverage and substantially more contigs than the other ATRun2 assemblies.

**Table 3:** Assembly statistics for ATRun2 SUP assemblies. Assemblies were evaluated against TAIR10. L50 is the minimum number of contigs needed to account for at least 50% of the total assembly length. The wtdbg2 assembly is shown before final polishing because polishing failed for this assembly.

| **Assembler** | **Contigs** | **Total length (Mb)** | **N50 (Mb)** | **L50** | **TAIR10 bases covered (%)** | **Average alignment identity (%)** |
|---|---|---|---|---|---|---|
| Canu | 63 | 122.44 | 12.62 | 5 | 99.84 | 99.96 |
| Flye | 49 | 119.52 | 13.59 | 5 | 99.68 | 99.96 |
| Miniasm | 122 | 118.69 | 2.21 | 20 | 96.45 | 99.95 |
| Pomoxis | 195 | 139.14 | 10.89 | 6 | 99.79 | 99.96 |
| Wtdbg2 | 1,353 | 151.95 | 2.47 | 14 | 97.45 | 99.84 |

**Fig 2.**
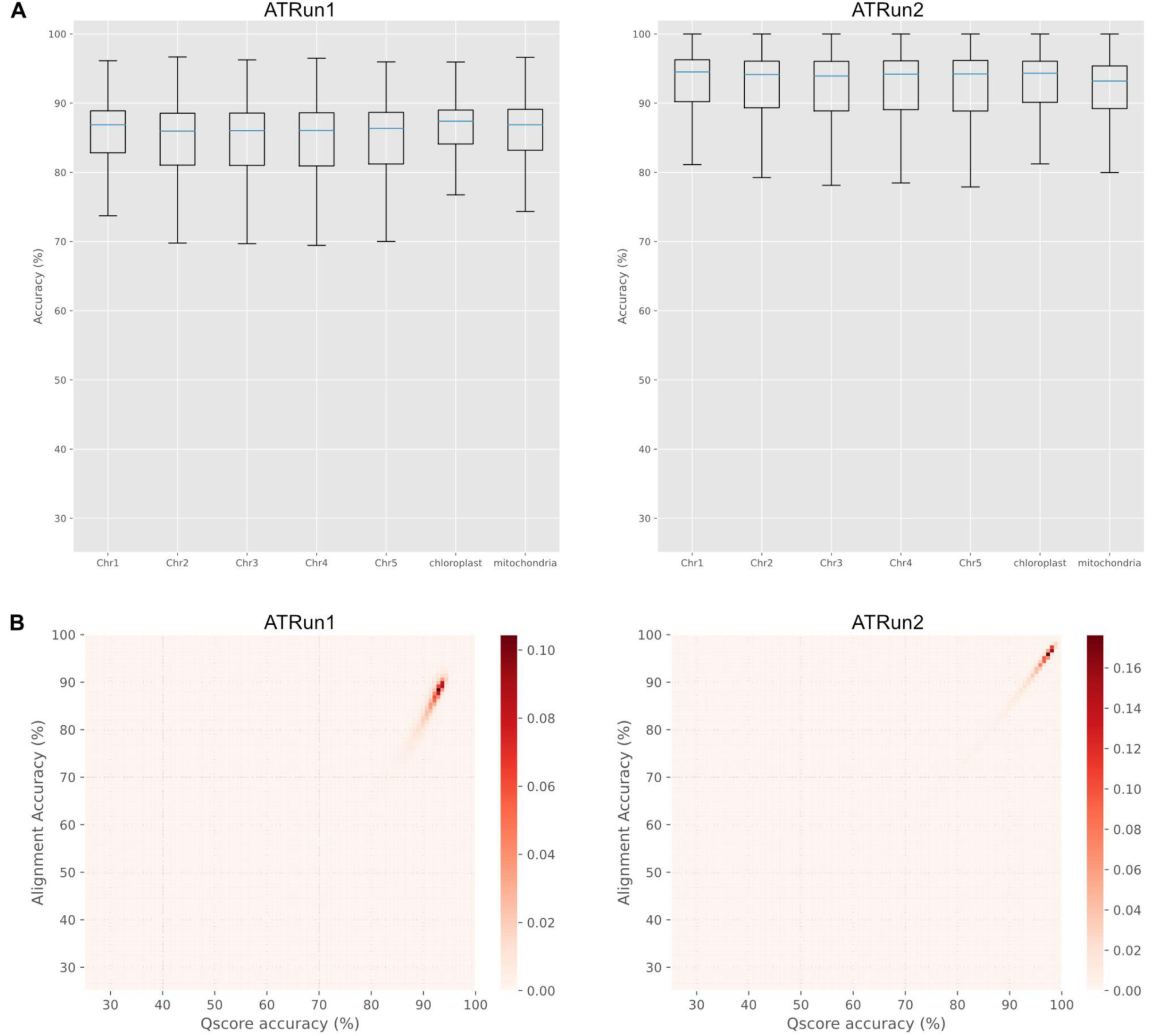
Alignment-based read accuracy for ATRun1 and ATRun2. **A.** Read alignment identity against TAIR10, grouped by reference chromosome or organellar sequence. **B.** Heatmaps comparing accuracy inferred from read Q scores with alignment identity for each run.

### Alignment-based read accuracy increased from R9.4.1 to R10.4

Alignment-based read accuracy increased from ATRun1 to ATRun2. Because both datasets were generated from the Col-0 accession, reads could be aligned to the TAIR10 reference to estimate read accuracy. ATRun2 reads had higher alignment-based accuracy than ATRun1 reads, and the ATRun2 accuracy distribution was narrower and shifted towards higher values (Fig 2A). Read Q scores were correlated with alignment-based accuracy for both datasets, although Q-score-derived accuracy was generally slightly higher than alignment-based accuracy, particularly for the older R9.4.1 ATRun1 dataset (Fig 2B).

### Current R10.4.1 data and long-read assembly produced a five-contig chromosome-scale nuclear assembly

To assess current nanopore assembly performance, we generated an additional R10.4.1 dataset, ATRun3, and processed it using recent basecalling software and a contemporary long-read assembly method. The rebasecalled ATRun3 dataset contained 706,072 reads with a total length of 14.30 Gbp, corresponding to approximately 106× coverage. The mean read length was 20.25 kbp and the read N50 was 37.54 kbp (Table 1).

Assembly of all ATRun3 reads without additional filtering produced an initial primary contig assembly of 199 contigs, with a total length of 160.39 Mbp, and an N50 of 26.15 Mbp. BUSCO analysis showed near-complete gene representation, with 99.9% complete BUSCOs [S:99.1%, D:0.9%], 0.1% fragmented, and 0.0% missing (Table 4). Although this assembly was highly complete at the gene level, it contained many smaller contigs, including low-coverage contigs and high-depth organellar sequences, indicating that curation was required to obtain a nuclear assembly.

**Table 4:** ATRun3 assembly curation and validation.

| **Assembly stage** | **Contigs** | **Total length (Mb)** | **N50 (Mb)** | **Query aligned to TAIR12 (%)** | **TAIR12 bases covered (%)** | **Average alignment identity (%)** | **Adjusted QV** | **BUSCO complete (%)** |
|---|---|---|---|---|---|---|---|---|
| Hifiasm primary contigs | 199 | 160.39 | 26.15 | 90.33 | 98.10 | 99.98 | 54.21 | 99.9 |
| Coverage-filtered (≥ 60) | 10 | 135.94 | 26.15 | 99.80 | 95.53 | 99.98 | 68.64 | 99.9 |
| Curated nuclear assembly | 5 | 134.93 | 26.15 | 99.99 | 95.53 | 99.98 | 68.85 | 99.9 |

Filtering contigs using hifiasm rd:i coverage tags with a threshold of rd:i ≥ 60 reduced the assembly to 10 contigs with a total length of 135.94 Mbp, while BUSCO completeness remained unchanged (Table 4). BLAST classification showed that the five smaller high-coverage contigs corresponded to organellar sequences, so these were excluded from the nuclear assembly. The final assembly consisted of five contigs with a total length of 134.93 Mbp and an N50 of 26.15 Mbp. BUSCO completeness was unchanged with 4,593 complete groups out of 4,596 assessed (99.9%), comprising 4,553 single-copy and 40 duplicated groups; three groups were fragmented and none were missing. The adjusted yak QV was 68.85, estimated from the same SUP reads used for assembly (Table 4). Whole-contig alignment to TAIR12 confirmed that the five contigs corresponded to the five A. thaliana nuclear chromosomes (Table 4; Fig 3). Comparison with TAIR12 using dnadiff gave 99.98% one-to-one alignment identity, with 99.99% of assembly bases aligned and 95.53% of reference bases covered. Together, these results show that 14.3 Gbp of current R10.4.1 nanopore data were sufficient to produce a highly accurate, chromosome-scale nuclear assembly of *A. thaliana* Col-0.

**Fig 3.**
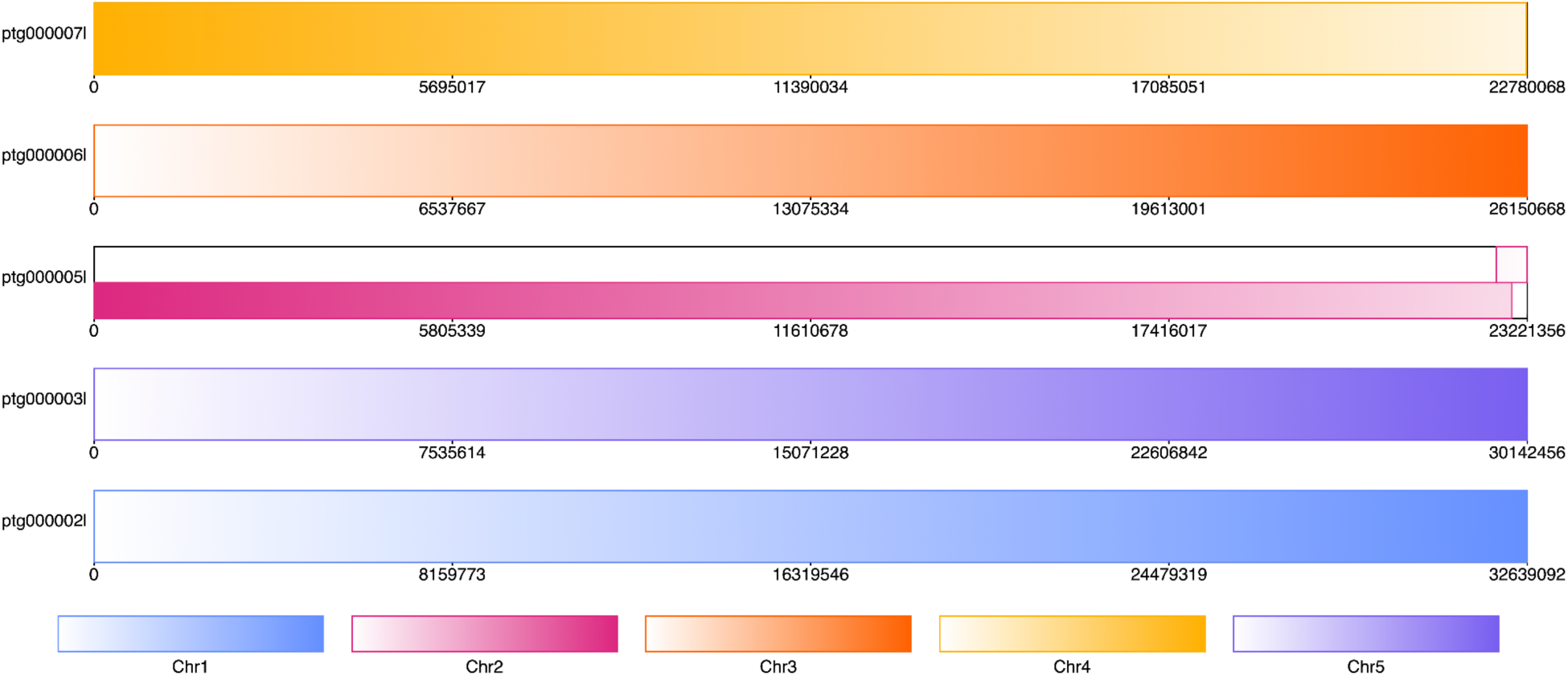
Whole-contig alignment of the curated ATRun3 nuclear assembly to TAIR12. The five retained ATRun3 contigs each align predominantly to one of the five *A. thaliana* nuclear chromosomes. Alignments shorter than 1% of the corresponding reference sequence were excluded from the visualisation.

### Nanopore CpG methylation calls correlated with bisulphite sequencing

To assess the utility of nanopore sequencing for epigenetic analysis, we extracted data on methylation at CpG sites from the R9.4.1 and R10.4 runs. We compared nanopore CpG methylation calls from ATRun1 and ATRun2 with a publicly available Illumina bisulphite sequencing dataset from *A. thaliana* Col-0 leaf tissue. Following coverage filtering and strand merging, nanopore and bisulphite methylation proportions were correlated at 461,613 shared CpG sites for ATRun1 (Pearson’s r = 0.93) and 1,144,748 shared CpG sites for ATRun2 (r = 0.96). The two comparisons shared 447,491 CpG sites.

**Fig 4.**
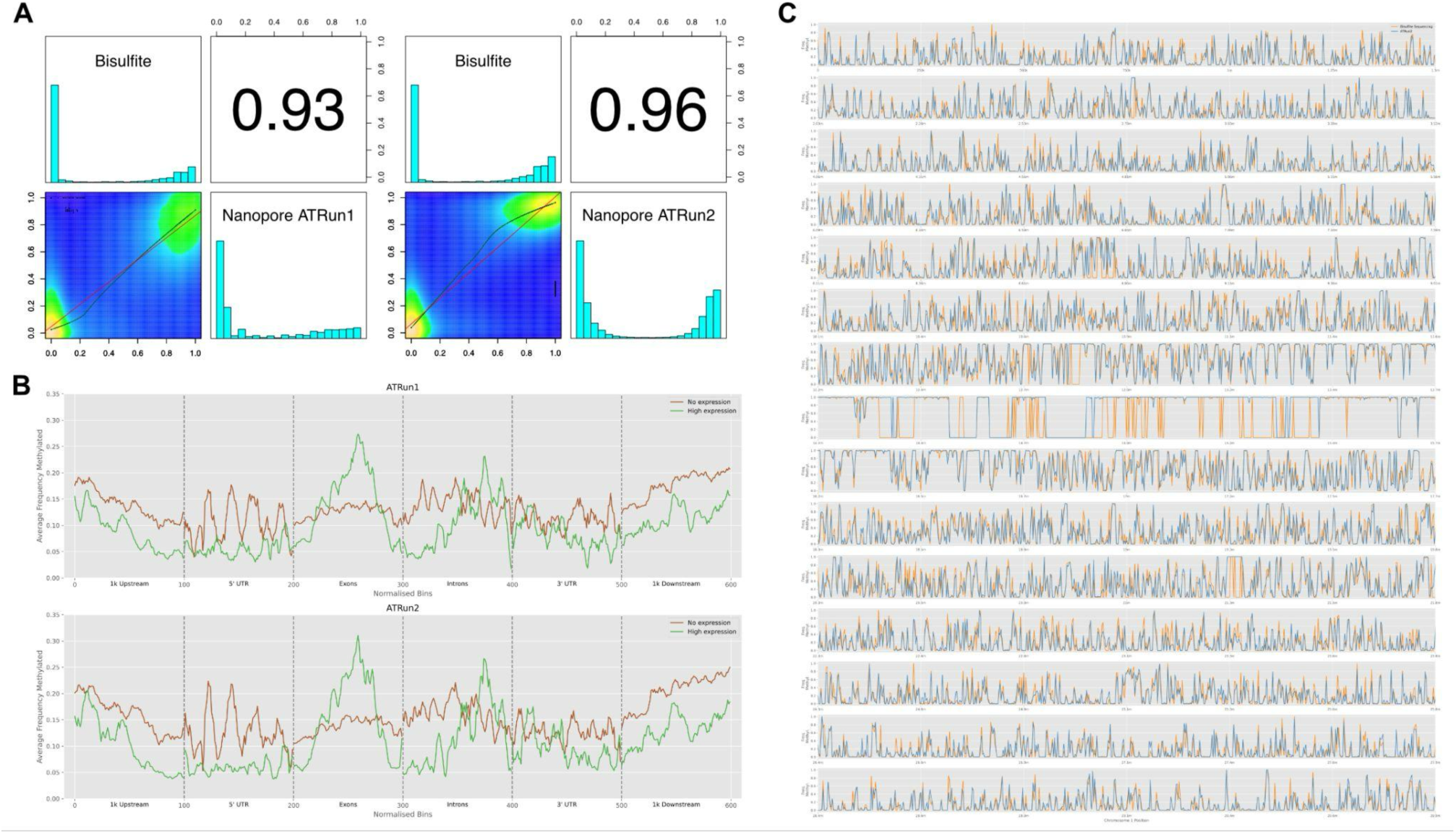
Comparison of nanopore and bisulphite CpG methylation estimates and gene-region methylation profiles. **A.** Pearson’s correlation coefficient (r) between CpG methylation proportions estimated using nanopore sequencing (ATRun1 and ATRun2) and bisulphite sequencing, calculated and plotted using the R package methylKit. Bar charts show the proportion of methylated reads at a particular site along the x-axis and the total proportion of such sites on the y-axis. **B.** Average gene-region CpG methylation profiles for highly expressed genes (n=1,001) and genes with no detectable expression (n=5,552) in leaf tissue. **C.** CpG methylation rate across Chromosome 1 using a tiling window of size 2.5 kbp. The blue track shows data from ATRun2, where only sites with coverage of at least 10 are included. The orange track shows data from Illumina bisulphite sequencing data, where only sites with coverage of at least 8 are included.

Chromosome-scale methylation profiles showed similar broad patterns between ATRun2 nanopore methylation calls and bisulphite sequencing across chromosome 1 (Fig 4C). These results indicate that the same nanopore datasets used for assembly can also recover CpG methylation patterns that are concordant with bisulphite sequencing at shared covered sites.

We next used nanopore CpG methylation calls to compare methylation profiles across gene bodies, including 5′ UTRs, exons, introns, and 3′ UTRs, and upstream and downstream flanking regions (Fig 4B). Genes were grouped according to leaf tissue expression data from Mergner et al. (2020), comprising 1,001 highly expressed genes (“High Expression”) and 5,552 genes with no detectable expression (“No Expression”).

Across both ATRun1 and ATRun2, genes with no detectable leaf expression showed higher average CpG methylation in upstream and downstream flanking regions, whereas highly expressed genes showed higher average CpG methylation across exon regions (Fig 4B). These patterns were consistent between the two datasets.

## Discussion

This study provides a practical comparison of nanopore-only *Arabidopsis thaliana* genome assembly and CpG methylation analysis across three datasets generated with different nanopore chemistries, basecalling models, and assembly workflows. Earlier R9.4.1 data were sufficient to generate multi-megabase and chromosome-arm-scale assemblies, but consensus accuracy and BUSCO completeness were limited when reads were basecalled with earlier models.

Rebasecalling the same raw signal data with a SUP model markedly improved read Q scores, consensus identity, and BUSCO completeness, showing that rebasecalling with improved basecalling models can recover additional value from existing nanopore signal datasets. The subsequent R10.4 dataset further improved read and assembly accuracy. Previous studies have demonstrated improved homopolymer resolution with R10.4 chemistry compared with R9.4.1 (Sereika et al., 2022), which may have contributed to the increased accuracy observed here. The R10.4.1/hifiasm workflow represented a further step change in assembly contiguity and consensus quality.

The ATRun3 assembly illustrates the current performance achievable with nanopore-only data for a compact plant genome. From 14.3 Gbp of R10.4.1 data, corresponding to approximately 106× coverage, hifiasm produced a highly complete assembly that could be curated to five nuclear contigs, each corresponding to one of the five *A. thaliana* chromosomes. The curated assembly had 99.9% complete BUSCOs, a non-independent adjusted yak QV estimate of 68.85, and 99.98% one-to-one alignment identity to TAIR12. This is a clear improvement over the chromosome-arm-scale assemblies generated from earlier R9.4.1 and R10.4 datasets in this study, reflecting both the increased accuracy of current nanopore reads and the development of long-read assemblers designed to make effective use of the data. Telomeric repeat arrays were detected at both ends of the contigs corresponding to chromosomes 1, 3 and 5, whereas the contigs corresponding to chromosomes 2 and 4 contained detectable arrays at only one end. Notably, the ATRun3 assembly was generated from simplex reads with a read N50 of 37.54 kbp, without specialised ultra-long-read library preparation. Although only 9.02% of reads exceeded 50 kbp, these contributed approximately 30× genome-equivalent coverage, with the longest read reaching 1.46 Mbp. This suggests that chromosome-scale assemblies, at least of compact plant genomes can be achieved without the challenge of generating exceptionally high read N50 values, although the contribution of the longest reads to assembly contiguity was not assessed independently.

The comparison between HAC and SUP basecalling of ATRun1 highlights the importance of retaining raw signal data. Rebasecalling the same R9.4.1 dataset improved mean read Q score and substantially increased BUSCO completeness across all assemblers, without changing the underlying biological sample or sequencing run. This suggests that archived nanopore signal data may remain useful as basecalling models improve, particularly for projects where resequencing is not practical. The effect was strongest for consensus accuracy and gene completeness, whereas contiguity changed less consistently, indicating that improved basecalling alone does not necessarily resolve all assembly graph or repeat-resolution limitations.

In addition to assembly, the same nanopore datasets identified CpG methylation patterns that were concordant with a published bisulphite sequencing dataset at shared sites passing coverage filters. This supports the use of nanopore data for combined assembly and methylation analysis and our results are broadly consistent with recent comparisons of long-read methylation detection and bisulphite-based methods. For example, Sigurpalsdottir et al. (2024) reported strong concordance between nanopore methylation calls and oxidative bisulphite sequencing in human DNA, using a per-CpG average Pearson correlation calculated across CpG sites. They also found that long-read sequencing detected more CpG sites than oxidative bisulphite sequencing. The stronger correlation observed for ATRun2 is consistent with improved methylation estimates, although differences in sequencing coverage, modification-calling models, and the sites retained for comparison may also contribute. As modified bases can be detected directly from the raw signal, nanopore sequencing avoids the additional sample treatment required for bisulphite sequencing and preserves long-range information that may be useful for studying methylation across repetitive regions, structural variants, and haplotypes.

The gene-region methylation profiles also produced expected biological patterns. Genes with no detectable expression in leaf tissue exhibited higher average CpG methylation in their flanking regions, whereas highly expressed genes showed greater CpG methylation across exons. The enrichment of CpG methylation within exons of expressed genes is consistent with gene-body methylation in plants, which is typically concentrated within transcribed regions and associated with transcriptional activity, although the relationship between methylation and expression is not necessarily linear (Zilberman et al., 2007). More broadly, the contrasting profiles of expressed and non-expressed genes are consistent with established associations between DNA methylation, gene architecture, and transcriptional state in plants (Law and Jacobsen, 2010). These results indicate that nanopore methylation calls can recover biologically meaningful CpG methylation patterns from the same sequencing datasets used for genome assembly. Although the present analysis focused on CpG methylation, nanopore-based methods have also been developed to detect methylation in the CHG and CHH contexts in plants, suggesting that broader epigenomic information may be recoverable from the sequencing data used for assembly (Ni et al., 2021).

Overall, our results show that nanopore-only sequencing can generate both genome assemblies and CpG methylation profiles for plant genomes, and that improvements in basecalling, sequencing chemistry, and assembly methods have substantially improved assembly quality over time. For *A. thaliana* Col-0, current R10.4.1 data combined with contemporary assembly software produced a five-contig nuclear assembly from a modest sequencing dataset, while the same nanopore platform also produced CpG methylation calls concordant with bisulphite sequencing at shared covered sites. The approach should be transferable to other eukaryotic organisms, but larger or more complex genomes may require higher sequencing depth, ultra-long-read preparation, phasing, scaffolding or additional curation, particularly where repeat content, heterozygosity, organellar DNA, or polyploidy complicate assembly. Rather than relying on manufacturer performance claims, this study provides an empirical example of how nanopore-only plant genome assembly and methylation profiling have improved across successive workflows.

## Acknowledgements

This research was supported in part by the NBI Research Computing through use of its high-performance computing cluster and associated data storage infrastructure.

## Author contributions

RML, MDC, and DH conceived and designed the study. DH, NP, NI, STH, and MG performed the experiments. SM, NP, YL, and RML analysed the data. RML, SM, NP, DH, and MDC wrote the manuscript. All authors read and approved the final manuscript.

## Data Availability

All raw read sequences presented in this work have been deposited in the European Nucleotide Archive under project accession PRJEB79853. Genome assemblies, analysis and plotting scripts, and supporting analysis data are available from Zenodo (https://doi.org/10.5281/zenodo.21628284).

---

**Table S1:** Effect of polishing on ATRun1 assembly statistics.

| **Assembly** | **Polished** | **Contigs** | **Total length (Mb)** | **N50 (Mb)** | **TAIR10 bases covered (%)** | **Average alignment identity (%)** |
|---|---|---|---|---|---|---|
| Canu | No | 128 | 120.6 | 10.66 | 99.73 | 98.52 |
| Canu | Yes | 128 | 121.89 | 10.76 | 99.74 | 98.88 |
| Flye | No | 106 | 120.72 | 14.09 | 99.83 | 98.97 |
| Flye | Yes | 90 | 120.89 | 14.12 | 99.83 | 98.92 |
| Pomoxis | No | 78 | 123.59 | 10.06 | 99.73 | 98.74 |
| Pomoxis | Yes | 78 | 123.35 | 10.07 | 99.72 | 99.22 |

**Table S2:** Effect of additional Racon polishing rounds on the ATRun1 Flye assembly. Both the one-round and four-round Racon treatments were followed by a final Nanopolish step.

| Assembly | Contigs | Total length (Mb) | N50 (Mb) | TAIR10 bases covered (%) | Average alignment identity (%) |
|---|---|---|---|---|---|
| Flye 1× Racon | 90 | 120.89 | 14.12 | 99.83 | 98.92 |
| Flye 4× Racon | 82 | 120.82 | 14.12 | 99.82 | 98.90 |

**Table S3:**
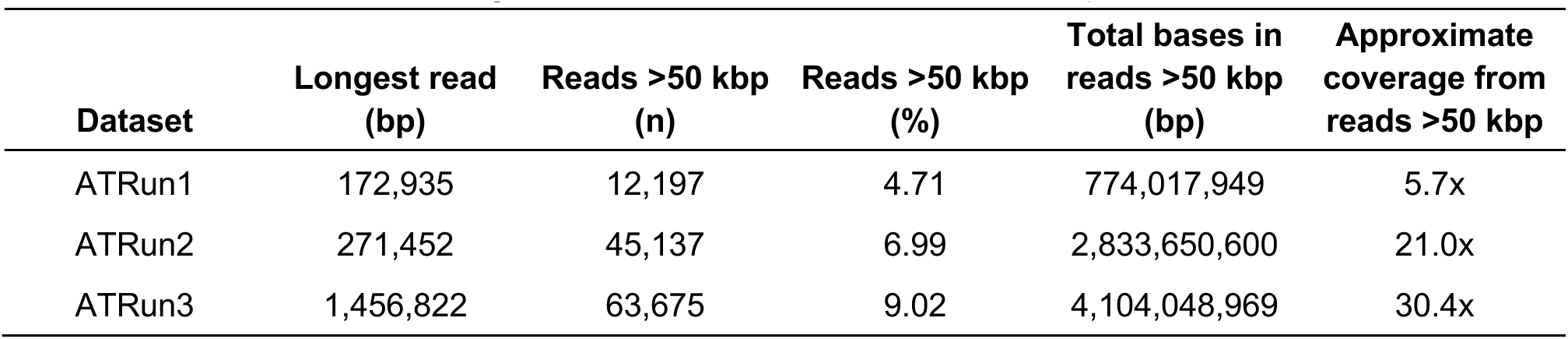
Long-read statistics for SUP-basecalled nanopore datasets.

